## Supplementary material for "Volumetric trans-scale imaging of massive quantity of heterogeneous cell populations in centimeter-wide tissue and embryo": Supplimental file

**This PDF file includes:**

Note 1-2
Figures S1-S6
Descriptions for Movie files (Movie S1-S9)

**Other supplementary materials:**
Movies S1–S9

**Note 1. Discussion on the confocal imaging system: Optical configuration, pinhole size and spatial resolution**

This section describes the details of the confocal system configuration and the setting of the pinhole size. AMATERAS in this paper is designed for fluorescence imaging in the visible wavelength region. Currently, the system is a single-channel observation system using excitation light at 470-490 nm and observing fluorescence in 510-540 nm region, which is primarily targeted at green-to-yellow fluorescent proteins such as EGFP (emission peak: 507 nm) and EYFP (emission peak: 527 nm). As the next step, we are planning to develop a two-channel observation system, which will also be capable of observing red fluorescent proteins, such as mRFP (emission peak: 607 nm) and mCherry (emission peak: 610 nm).

<u>Excitation system</u>

The point spread function (PSF) of a confocal microscope is expressed by multiplying the distribution of the focused spot of the excitation light by the PSF of the imaging system of fluorescence. In the excitation system, the pinhole is illuminated by the LED light (center wavelength: 470 nm), and the light is diffracted by the pinhole, causing the light to spread at the tube lens side (Fig. S2A). The diffracted light is collected and collimated by the tube lens and focused into the sample by the objective lens. The diffraction of light through a circular aperture is expressed by a well-known formula involving the Bessel function,

$$U(\theta) \propto \frac{J_1(ka\sin\theta)}{ka\sin\theta} \tag{1}$$

where $J_1(u)$, $k$, $a$, $\theta$ denote the Bessel function of the first kind with the order of 1, wavenumber, aperture radius, and diffraction angle, respectively. The angular dependence of intensity can be obtained by taking a square of Eq. (1) (Fig. S2B). The pinhole size in the current setup is 6 μm, at which the diffraction pattern has the first dark ring at about 0.1 radian (Fig. S2B). Since our tube lens has an NA of 0.125, the diffracted light does not fill the entire NA of the lens, and as a result, the effective NA of the focused light into the sample is lower than 0.25 (Fig. S2A). This differs from a typical, laser-based, single-focus scanning confocal microscope. On the other hand, the current configuration is an appropriate choice in terms of light efficiency since most of the diffracted light enters within the NA of the tube lens.

Fluorescence imaging system

As for the fluorescence imaging system (Fig. S2C), fluorescence light emitted from fluorescent molecules in the sample is collected by the objective lens with 0.25 NA. Since the fluorescence is emitted isotropically, it fills the entire object-side NA and is focused onto the pinhole by the tube lens with 0.125 NA. If the pinhole diameter is small enough (mathematically represented by the delta function), the focal light field spot focused by the objective lens at the fluorescence wavelength can be considered to correspond to the PSF of the imaging system. Figure S2D shows the ideal PSF. This PSF was obtained by numerical calculation of the scalar wave integration.

Selection of the pinhole diameter

Here, the selection of the pinhole size (diameter) for our optical system is discussed. The pinhole diameter of a confocal microscope is usually set based on the Airy disk diameter of the spot of fluorescence focused on the pinhole. In general-purpose instruments, the pinhole diameter is set to about the Airy disk diameter, but when high spatial resolution is required, it is set to a value smaller than the disk, *e.g.*, half the diameter of Airy disk. At the green wavelength region (*e.g.*, EGFP, EYFP) and the red wavelength region (*e.g.*, mRFP, mCherry), which are currently considered as targets for AMATERAS observation, the Airy disk diameters of the spots focused by our tube lenses (NA = 0.125) are about 5 μm (EGFP: 4.95 μm, EYFP: 5.14 μm) and 6 μm (mRFP: 5.92 μm, mCherry: 5.95 μm), respectively, where we used the following equation for the Airy disk diameter, where $\lambda$ denotes the wavelength.

$$d_{Airy} = 1.22\frac{\lambda}{NA} \tag{2}$$

The design of the pinhole diameter was considered for both of these systems, excitation and fluorescence imaging. In our optical system, the transmittance of the pinhole disk is low, so it is not desirable to make it smaller than the Airy disk to further reduce its efficiency. On the other hand, if the pinhole is made too much larger than the Airy disk, it is meaningless because it will not provide 3D resolution. Considering that it can be used in both of the two wavelength ranges, we decided on a pinhole size of 6 μm, which is comparable to the Airy disk diameter in the red wavelength range.

Theoretical and actual resolution in the longitudinal direction

The theoretical resolution (FWHM) in the $z$-direction in the ideal confocal fluorescence microscope optical system (excitation light is focused to the diffraction limit and detected by a detector of infinitesimal size) is approximately expressed by the following equation with NA, the wavelength of excitation light ($\lambda_{ex}$) and refractive index of immersion medium ($n$).

$$\Delta z_{cf} = 0.64 \frac{\lambda_{ex}}{n - \sqrt{n^2 - NA^2}} \quad (3)$$

The value of $\Delta z_{cf}$ for $NA = 0.25$, $\lambda_{ex} = 470$, and $n = 1$ is 9.5 μm, which is considered the best possible resolution that can be achieved with our lens system. As mentioned above, the focusing NA of the excitation light is smaller than the NA of the objective lens, and the pinhole size is slightly larger than the Airy disk. These factors limit the resolution to more than 13 μm at present (Fig. 3D-E). The challenge for the future is to resolve these issues and approach the ideal resolution.

Theoretical and actual resolution in the transverse direction

The theoretical spatial resolution (FWHM) in the transverse direction in the ideal confocal fluorescence microscope system is approximately expressed by the following equation.

$$\Delta x_{cf} = 0.37 \frac{\lambda_{ex}}{NA} \quad (4)$$

The value of $\Delta x_{cf}$ for $NA = 0.25$ and $\lambda_{ex} = 470$ is 0.70 μm. However, the experimental value (Fig. 3D-E) is significantly larger than this theoretical value. Similar to the above discussion on the resolution in the $z$-direction, this is attributed to the fact that our imaging system is far from the ideal system. In addition, the resolution in the transverse direction is also influenced by the secondary relay-lens system that transfers the fluorescence light component passing through the pinhole to the image sensor with magnification of 1× or 2× (Fig. 3B). Since the NA of the 1× relay lens (0.079) is significantly smaller than that of the tube lens (0.125), large NA component (> 0.079) is lost at the edge of the entrance aperture of the relay lens. The effective NA for the fluorescence imaging is 0.158 at the sample space, which is the NA of the relay lens multiplied by the magnification factor (2×). This is the reason why the spatial resolution of AMATERAS-2c in the transverse direction (Fig. 3D) is downgraded from the wide-field imaging system (AMATERAS-2w) (Fig. 2A) as well as from the ideal confocal imaging system. The transverse resolution (FWHM) of the theoretical PSF for wide-field fluorescence imaging is expressed by the following equation with NA and emission wavelength ($\lambda_{em}$).

$$\Delta x_{wf} = 0.51 \frac{\lambda_{em}}{NA} \quad (5)$$

The values of $\Delta x_{wf}$ for $NA$ = 0.25 and 0.158 are 1.05 μm and 1.66 μm ($\lambda_{em}$ = 515 nm), respectively. This difference explains the degradation of the resultant transverse resolution.

In the case when the 2× relay lens is used, on the other hand, its NA (0.12) is only slightly smaller than that of the tube lens (0.125). The degradation of the transverse resolution is well mitigated compared to the 1× relay lens (Fig. 3E).

### Note 2. Discussion on the computational sectioning: Parameter adjustment and limitation

Basic concept

In the optimization of the cutoff frequency of the baseline estimation, we applied the concept of independent component analysis (ICA). As similar to ICA, we assumed that fluorescence image in the focal plane is away from a Gaussian distribution (normal distribution), but the superposition of images from multiple planes including in-focus and out-of-focus planes brings a distribution closer to a Gaussian distribution. Figure S4 shows how the estimated image varies depending on the cutoff frequency. The raw image (Fig. S4A) is a part of the cardiac organoid shown in Fig. 4 in the main text. The computational sectioning calculation was applied with various cutoff frequencies ($F_c$) from 0.010 to 0.300 $\mu m^{-1}$, and the four types of statistical measures to represent the non-Gaussian nature (non-Gaussianity) were calculated for the estimated in-focus images. The non-Gaussianity measures are kurtosis, skewness, negentropy, and mutual information (48). The in-focus and out-of-focus images at three cutoff frequencies ($F_c$ = 0.020, 0.063, 0.141 $\mu m^{-1}$) are shown to see how the images change with the frequency (Fig. S4B-C), and the line profiles on the raw image, in-focus, and out-of-focus images are shown in Fig. S4D to clarify their difference.

Determination of the non-Gaussianity measure and the cutoff frequency

First, we decided which measure we should use for the non-Gaussianity. Figure S4E shows the cutoff frequency dependence of the four measures, which was calculated for the subregion of image (yellow square region in Fig. S4A). In ICA, maximizing kurtosis or skewness is considered to maximize the non-Gaussianity, whereas minimizing negentropy or mutual information is considered to maximize the non-Gaussianity. Among these measures, negentropy and mutual information have their minimum around 0.141, but the estimated baseline contains the nuclear structure. The in-focus image, obtained by subtracting the baseline image from the original image, is noise-dominant (Fig. S4D, right). This is not suitable for our purposes, so we do not employ these indices. On the other hand, the skewness has a maximum value around 0.063, where the in-focus image appears to have clear nuclei, which is consistent with our objective (Fig. S4B, middle). When the frequency was much lower ($F_c$ = 0.020) (Fig. S4B, left), the background light component remained in the in-focus image. The kurtosis was found to have a similar frequency response to the skewness, with its maximum value at almost the same frequency. This trend was almost common in other images as well. These discussions led us to adopt skewness (or kurtosis) as non-Gaussianity measures, and set the cutoff frequency to $F_c$ = 0.063 for this image data, the frequency with the maximum skewness value. This optimization of the cutoff frequency was performed on a subset of a given data prior to the calculation of the entire dataset.

<u>Limitation of the computational sectioning</u>

Here, we discuss the limitation of applicability of our computational sectioning. As this method involves a baseline estimation process using a low-pass filter with a single cutoff frequency, the structures of the target object should be of uniform size, such as the cellular nucleus. When different sizes are mixed together in the object, the image processing often does not work well. An example of the application of this method to a computer-generated model image is shown in Fig. S5. In the model, a fluorescent sphere with a diameter of 10 μm or 30 μm is centered as the target object, and a 50 μm fluorescent sphere is placed 100 μm deep as the source of background light. A pseudo fluorescence image was generated on that 3D fluorescence distribution by the following procedure: (1) A theoretical point-spread function of the wide-field imaging system (NA = 0.25, wavelength = 520 nm) was convolved, (2) the intensity values was adjusted so that the maximum fluorescence intensity was 100, and (3) the values were converted to integers by rounding and Poisson noise was added. Figure S5A-top shows a fluorescence image of the *z*-plane passing through the center of a 10 μm target sphere. The in-focus image was estimated (Figure S5A-bottom) by the method described above, with the optimal cutoff frequency set to the frequency at which the skewness is maximum, $F_c$ = 0.045 (Fig. S5C). Similarly, Figure S5B shows the original image of a 30 μm sphere and the estimated in-focus image. For the 30 μm sphere, a cutoff frequency of 0.022 was found to be optimal (Fig. S5C), with which the in-focus image was estimated (Fig. S5B, bottom left). However, an artifact occurred where the area outside the boundary formed a valley, which happened because the spatial frequencies of the background light and the target sphere were close, leading to insufficient separation. If the cutoff frequency chosen for the 10 μm sphere is used for the 30 μm sphere image, the image of the target sphere falls within the baseline (Fig. S5B, bottom right). Figure S5D shows images estimated using several frequencies in the case of 10 μm and 30 μm spheres side by side. At $F_c = 0.045$, the optimal frequency for 10 μm sphere (Fig. S5C), the 10 μm sphere is still well separated from the background, but the 30 μm sphere is included in the background light. Many artifacts occur at other frequencies as well. These results indicate that it is undesirable to apply this method to fluorescent images of two objects of very different sizes or of objects that are close in size to the background light.

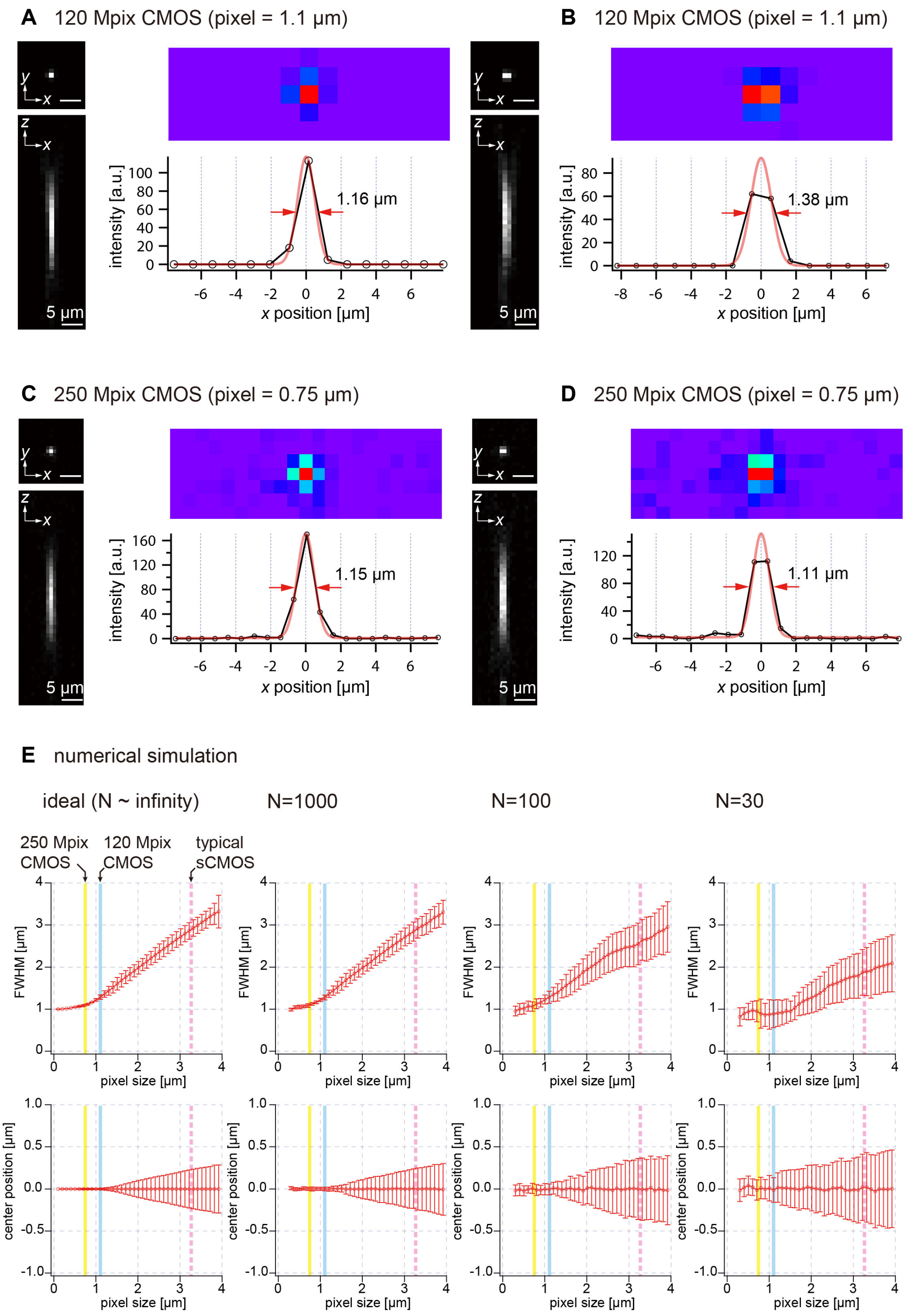


**Fig. S1.**

Uncertainty in spatial resolution and position owing to undersampling. (A),(B) Two cases of images of fluorescent beads (diameter, 0.2 μm) captured using the 120-megapixel camera. (C),(D) Two cases of images of fluorescent beads (diameter, 0.2 μm) captured using the 250-

megapixel camera. (E) Dependence of FWHM and center position on pixel size, as revealed by a numerical simulation. The number of photon ($N$) is varied from $N \sim$ infinity, 1000, 100, and 30.

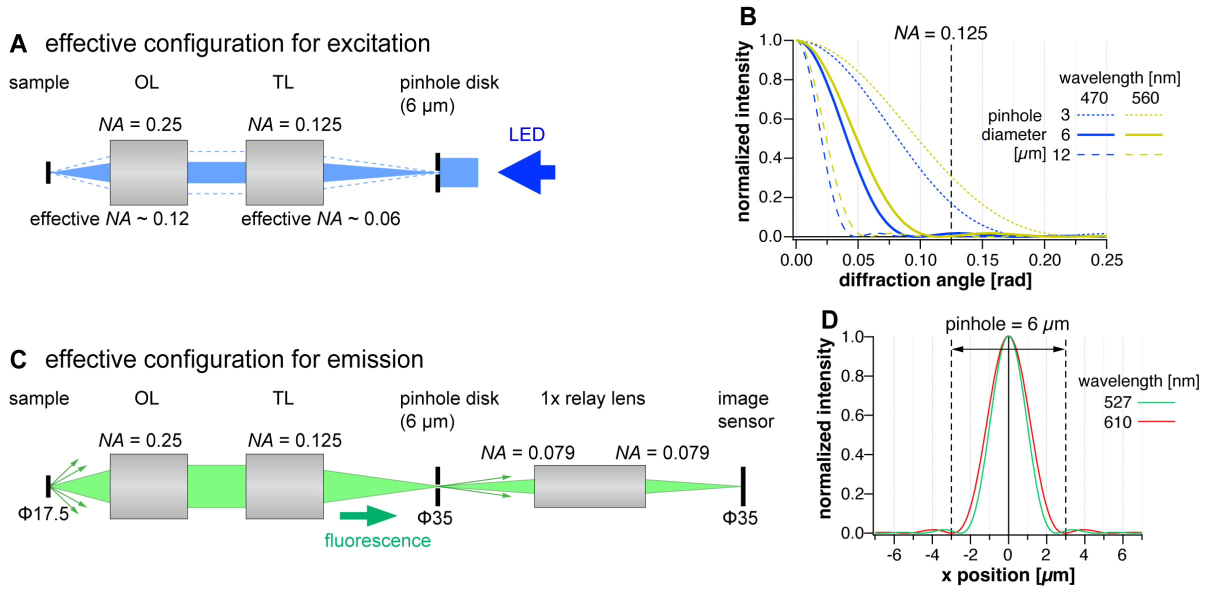


**Fig. S2.**

(A) Effective configuration for excitation in the confocal imaging system, AMATERAS-2c. OL and TL denote objective lens and tube lens, respectively. (B) Diffraction pattern by a pinhole with various conditions (wavelength and pinhole diameter), calculated by Eq. 1. The intensity of diffracted light is plotted as a function of diffraction angle. The diffraction angle equivalent to the NA of the tube lens is represented by the dotted line. (C) Effective configuration for emission in AMATERAS-2c. (D) Intensity distribution of focused light spot by tube lens with 0.125 NA at the two fluorescence wavelengths (527 and 610 nm). The size of the 6 μm pinhole is indicated by the dashed lines.

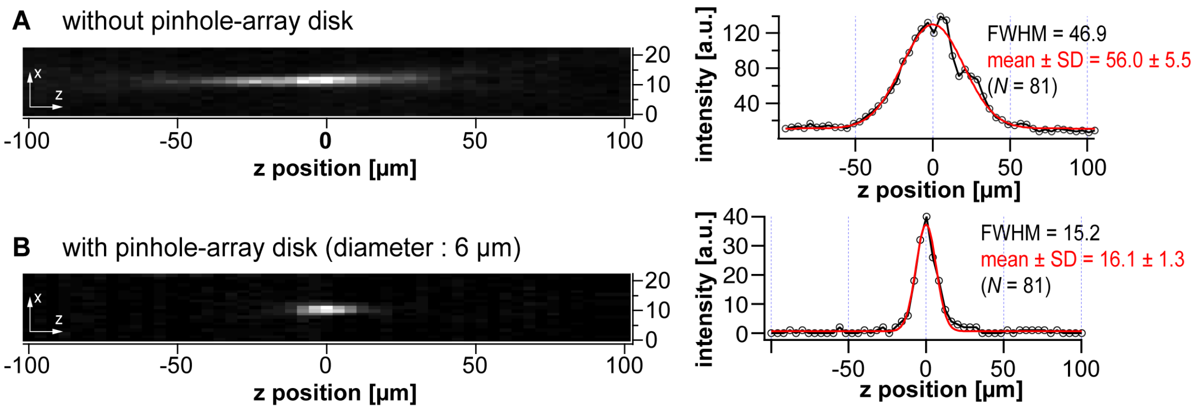


**Fig. S3.**

(A-B) Effect of confocality evaluated by point-spread function measured in the absence (A) and presence (B) of the pinhole-array disk (diameter: 6 µm). The optical configuration for this experiment employed the 1× relay lens, the same as used in Fig. 3D in the main text (total magnification: 2×). Left panels show *xz* planes centering at single fluorescent bead. Right panels show the line profiles in the *z* direction across the bead positions in the left panels. The FWHMs are noted alongside the line profiles. The statistical values of 81 peaks are also shown as well.

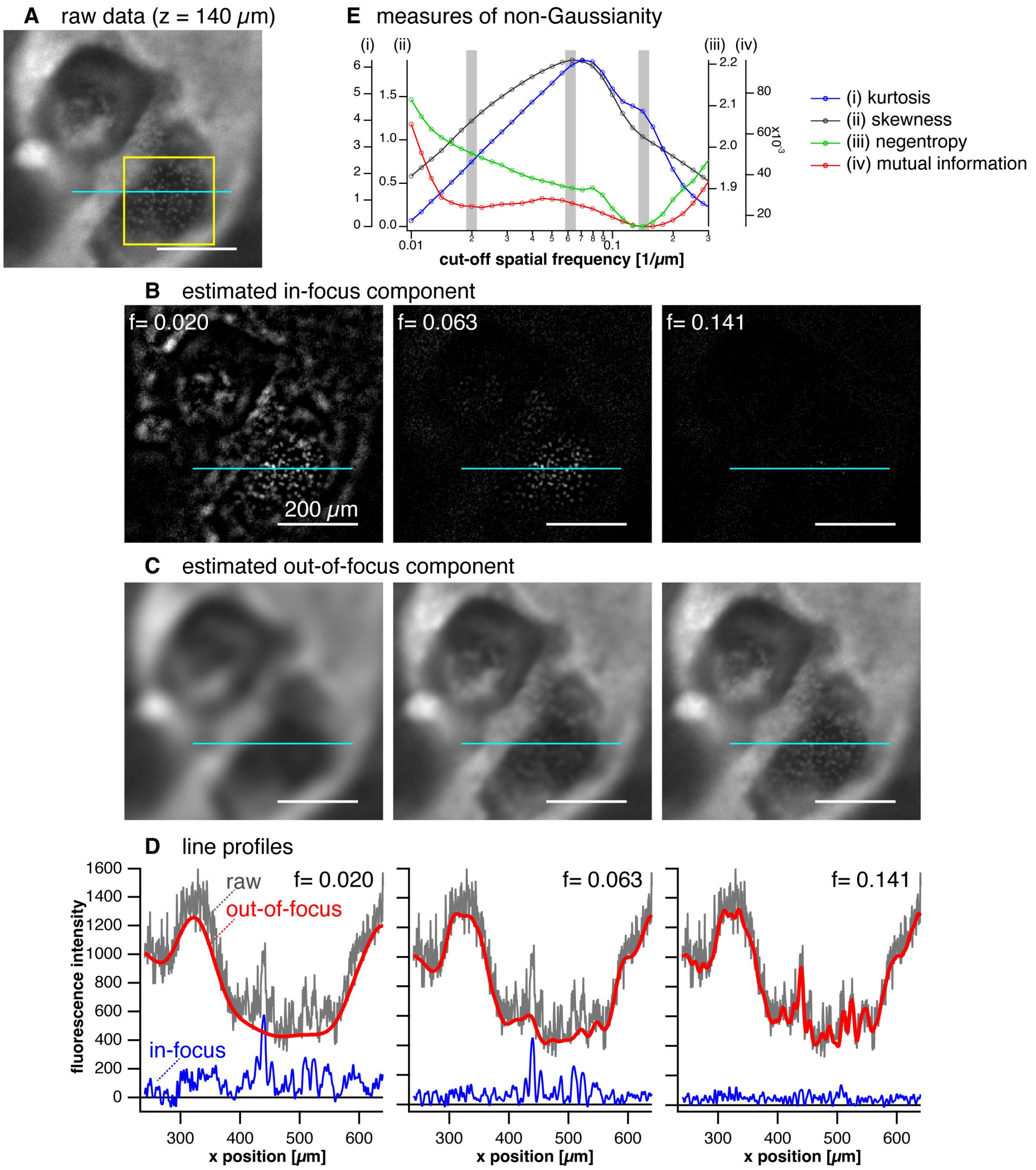


**Fig. S4.**

Performance of computational sectioning with varied cutoff frequency. (A) A raw image data, a local area of a *z*-layer (*z* = 140 μm) in a *z*-stack data of a dome structure of a myocardial organoid, which is shown in Fig. 4 in the main text. (B-C) Estimated in-focus images (B) and out-of-focus images (C) for three different cutoff frequencies, $F_c$ = 0.020, 0.063, and 0.141. Scale bar (white line): 200 μm. (D) Line profiles of intensity on the light-blue lines drawn in A, B, and C. (E) Four types of statistical measures for non-Gaussianity calculated as a function of cutoff frequency of the low-pass filter. The values were calculated for the in-focus image in the yellow square region. The three frequencies used for the image examples (B-D) are indicated by gray bars.

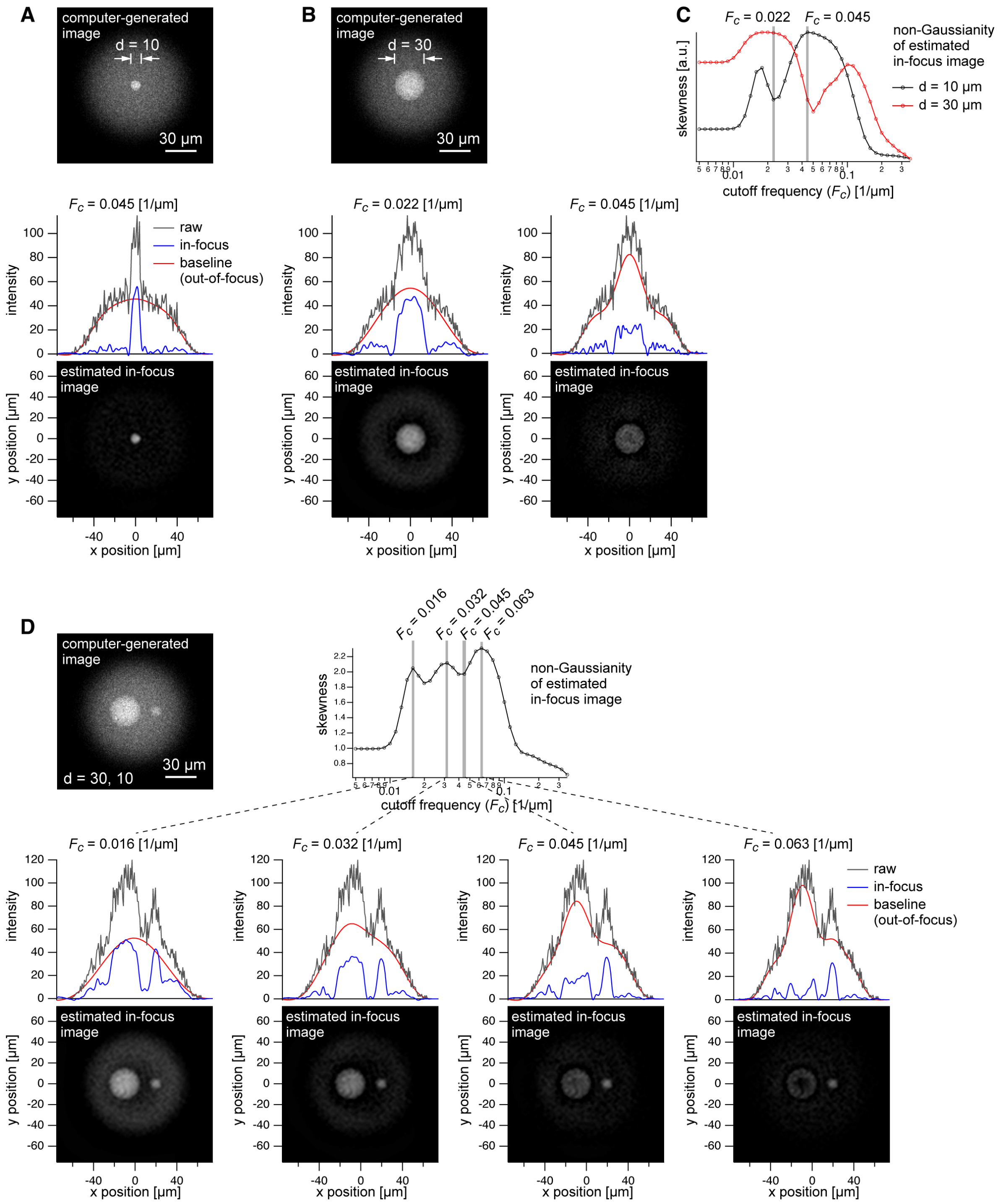


**Fig. S5. Dependence of baseline estimation performance on object size**

(A-B) Top: Computer-generated fluorescence image of superposition of a 10 μm sphere and a 30 μm sphere (object) and a 50 μm sphere placed 100 μm deep (background). Bottom: Estimated in-focus image at the cutoff frequency of 0.045 and 0.022 (only in B). Line profiles of the raw, estimated in-focus and out-of-focus images are shown on the images. (C)

Frequency dependence of the non-Gaussianity (skewness) of in-focus image for the 10 µm and 30 µm spheres. (D) Top: Computer-generated fluorescence image of superposition of a 30 µm sphere and a 10 µm sphere (object) and a 50 µm sphere placed 100 µm deep (background). Bottom: Estimated in-focus image at four cutoff frequencies (0.016, 0.032, 0.045, and 0.063) shown with line profiles of raw, in-focus and out-of-focus images.

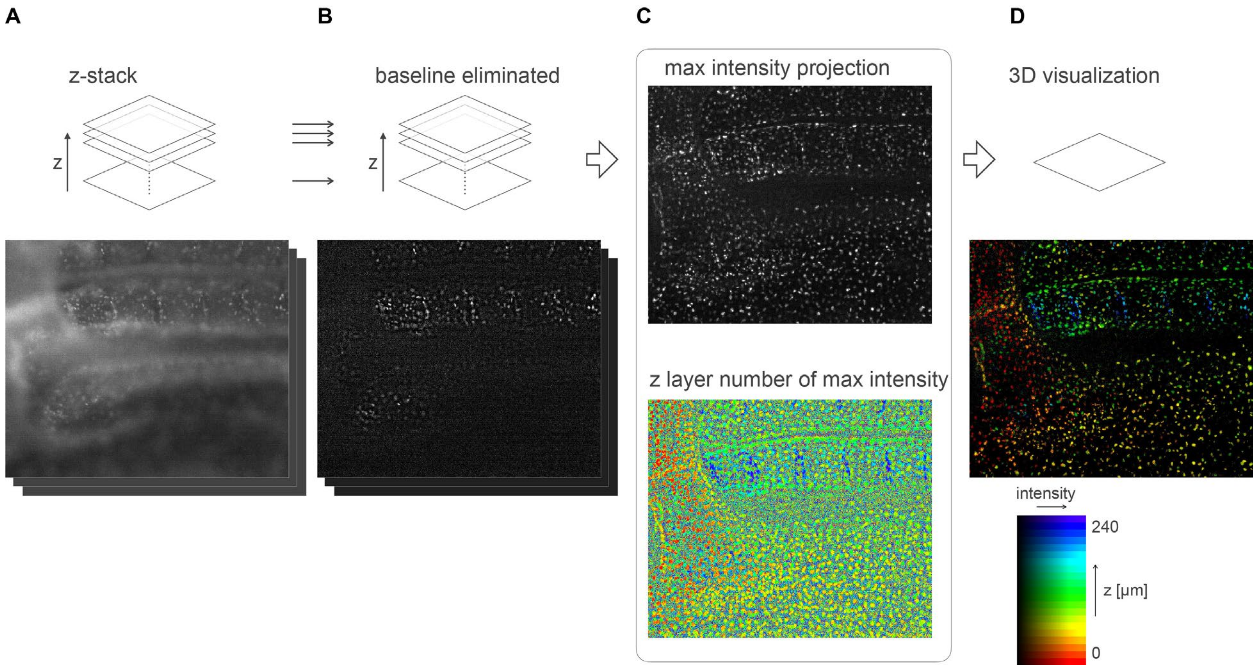


**Fig. S6.**

Demonstration of computational processing and 3D visualization of the *z*-stack data of quail embryo. (A) Raw image data. (B) In-focus component obtained by computational sectioning to eliminate the baseline component (out-of-focus component). (C) Images of projected max intensity and the z-layer number where maximum intensity is found, which are calculated from a set of *z*-stack. (D) Visualization of the 3D data using the HSB color table.

**List of supplementary movie files**

**Movie S1.**

A *z*-stack of a cavity chamber structure of myocardial organoid (left) without and (right) with the computational sectioning, corresponding to Figs. 4C and 4D, respectively. The *z*-position is swept with a step of 4 μm in 56 layers. Magenta and cyan colors represent the distribution of immunostained cardiac troponin T and Hoechst-labeled nuclei, respectively.

**Movie S2.**

A 3D isosurface representation of a dome structure of cavity chamber with varying the view angle. The cropped region is indicated by a dashed square in Fig. 4D.

**Movie S3.**

A *z*-stack of the mouse brain section shown in the full coronal plain region (Fig. 5B). The z-position is swept with a step of 4 μm in 378 layers.

**Movie S4.**

A *z*-stack of the mouse brain section shown in the local volume corresponding to Fig. 5C. The *z*-position is swept with a step of 4 μm in 378 layers

**Movie S5.**

A *z*-stack of the mouse brain section shown in the local volume in the cortex region. Cells detected by ELEPHANT are indicated by green oval markers. The *z*-position is swept with a step of 4 μm in 50 layers (200 μm) from the surface of the brain section.

**Movies S6–S8.**

Time lapse movie of the quail development in: (S5) the entire body region, (S6) ventral aorta region, and (S7) dorsal aorta region. The frame interval is 7.5 min in 25 h (200 frames). The rainbow color of cells denotes the *z*-position based on the color scale shown in Fig. 6C.

**Movie S9.**

Time lapse movie of the dorsal aorta region (Fig. 6E) in the 3D isosurface representation.
